# Molecular signatures of selection associated with host-plant differences in *Pieris* butterflies

**DOI:** 10.1101/627182

**Authors:** Yu Okamura, Ai Sato, Natsumi Tsuzuki, Masashi Murakami, Hanna Heidel-Fischer, Heiko Vogel

## Abstract

Adaptive traits that enable organisms to conquer novel niches and experience subsequent diversification are ecologically and evolutionarily important. The larvae of *Pieris* butterflies express nitrile-specifier proteins (NSPs), a key innovation for overcoming the glucosinolate (GLS)-myrosinase-based defense system of their Brassicales host-plants. NSPs are a member of the NSP-like gene family, which includes the major allergen (MA) protein, a paralog of NSP with a GLS-disarming function, and a single domain major allergen (SDMA) protein, whose function is unknown. The arms-race between a highly variable host-plant defense system and members of the NSP-like gene family is suggested to mediate diversification in both Pierid butterflies and Brassicales plants. Here, we combined feeding experiments using 25 Brassicaceae plants and five *Pieris* species with larval transcriptome data to investigate the evolutionary forces acting on NSP-like gene family members associated with patterns of host-plant usage. Although we observed significantly elevated nonsynonymous to synonymous substitution ratios in NSPs, no such pattern was observed in MAs or SDMAs. Furthermore, we found a signature of positive selection of NSP at a phylogenetic branch which reflects different host-plant preferences. Our data indicate that NSPs have evolved in response to shifting preferences for host plants among five *Pieris* butterflies, whereas MAs and SDMAs appear to have more conserved functions. Our results show that the evolution and functional differentiation of key genes used in host-plant adaptation play a crucial role in the chemical arms-race between *Pieris* butterflies and their Brassicales host-plants.

## Introduction

Key innovations that enable organisms to acquire novel niches and experience subsequent radiation are ecologically and evolutionarily important (Bond & Opell, 1988; Hunter, 1998). In plant-herbivore interactions, a number of key innovations were identified that enabled herbivores to overcome specific plant defense mechanisms and colonize novel host-plants (Berenbaum, Favret, & Schuler, 1996; Janz, 2011; Wheat et al., 2007). *Pieris* butterfly larvae use plants containing glucosinolate (GLSs) as hosts, redirecting toxic breakdown products to less toxic metabolites using gut-expressed nitrile-specifier proteins (NSPs) (Wittstock et al., 2004). NSPs are known to be a key innovation of *Pieris* butterflies: the acquisition of NSPs enabled *Pieri*s to colonize GLS-containing Brassicales followed by higher speciation rates compared to those of sister butterfly clades (Edger et al., 2015; Fischer, Wheat, Heckel, & Vogel, 2008; Heidel-Fischer, Vogel, Heckel, & Wheat, 2010; Wheat et al., 2007).

NSPs are members of the small NSP-like gene family, which includes major allergen (MA) proteins and single domain major allergen (SDMA) proteins (Fischer et al., 2008). The functions of MAs and SDMAs are mostly unclear, however, the structures of MAs and NSPs are known to be similar: three replicated domains originated from SDMAs (Fischer et al., 2008). In addition, although SDMAs are generally expressed in the guts of Lepidopteran larvae (Randall, Perera, London, & Mueller, 2013), NSP and MA are only found in Pierid butterflies feeding on Brassicales (Fischer et al., 2008). These findings suggest that in *Pieris*, MAs, as in NSPs, have a function related to disarming GLSs. The ability of MAs to redirect GLS hydrolysis was recently documented in one Brassicales-feeding Pierid, *Anthocharis cardamines*, which seems to have MA genes only, that is, it lacks NSPs (Edger et al., 2015). Thus, although the function of MAs in Pieridae is largely unknown, especially in those species which have NSPs and MAs, MAs also appear to be ecologically important for overcoming the host plant’s GLS-based defense system.

Previous studies indicated that the co-evolutionary diversification of Brassicales plants and Pierid butterflies was mediated by the chemical arms-race between the glucosinolate-myrosinase defense system and members of the NSP-like gene family (Edger et al., 2015). Past increases of GLS complexity in Brassicales were followed by the evolution in Pierid butterflies of NSP-like gene family members, suggesting that members of the NSP-like gene family would potentially be under strong selection pressure, were Pieridae butterflies to expand or shift their host plants. Such a scenario is supported by recent findings of signatures of positive selection in partial NSP sequences of a pair of *Pieris* butterflies in comparison with the signatures of 70 randomly selected genes (Heidel-Fischer et al., 2010). However, the evolutionary forces acting on all NSP-like gene family members, especially when considering the associated host plant spectrum, remains unknown.

Here, we focus on five Japanese butterfly species (*Pieris napi*, *P. melete*, *P. rapae*, *P. brassicae* and *P. canidia*) in the genus *Pieris*, which has both NSP and MA genes, and feed on Brassicaceae plants with the highest GLS diversity among the Brassicales. The five *Pieris* species have different host spectra according to field observations (Fig. 1), with *P. napi* and *P. melete* frequently using wild Brassicaceae plants (such as *Arabis* or *Arabidopsis*), whereas *P. rapae* and *P. brassicae* tend to feed on Brassicaceae crops and are known as major pests (Benson, Pasquale, Van Driesche, & Elkinton, 2003; Kitahara, 2016; Ohsaki & Sato, 1994; Ueno, 1997). In contrast, in Japan, *P. canidia* can be found only in the southern islands (Yonaguni Island, Okinawa), relying on the limited number of host plants, such as *Cardamine* or *Lepidium*, in their habitat range.

**Fig. 1.**
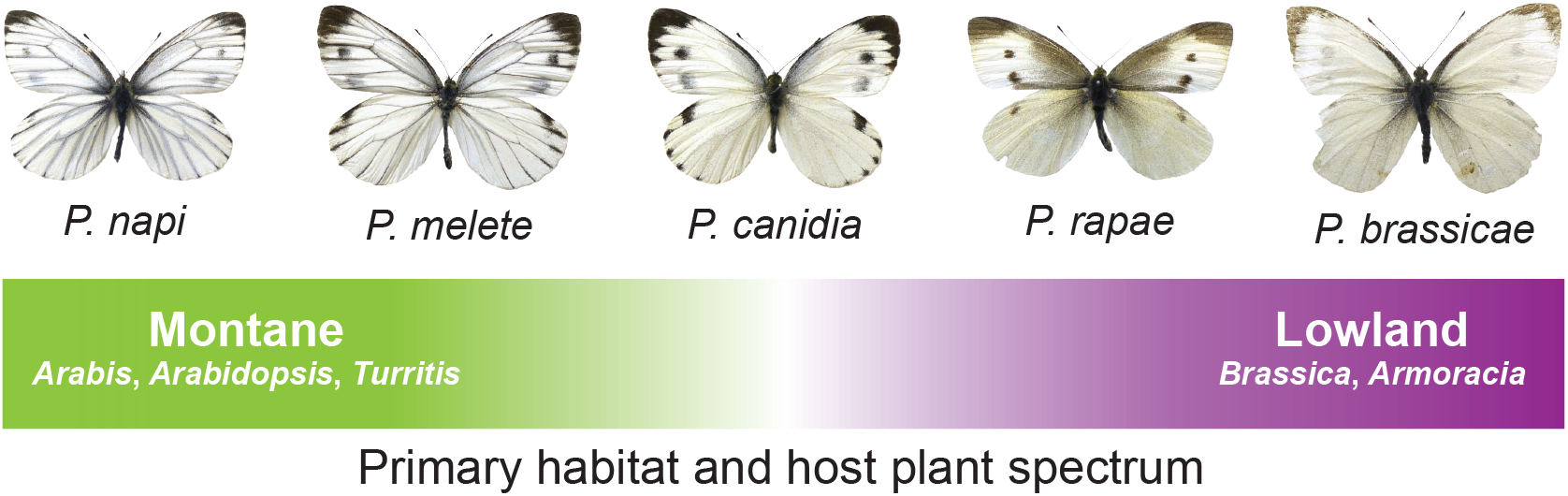
Field observations of primary habitat and larval host-plant spectra of five *Pieris* butterflies in Japan. *P. napi* and *P. melete* tend to be found in montane habitat and rely mostly on Brassicaceae plants in forests; these include *Arabis*, *Arabidopsis* or *Turritis*. *P. rapae* and *P. brassicae* are known as Brassica crop pests. In Japan, *P. canidia* can only be found in a restricted area and uses *Cardamine* or *Lepidium* as host plants.

With larval transcriptome (RNA-seq) data from the five *Pieris* species, we analyzed the divergence in amino acid sequences based on nonsynonymous (dN) and synonymous substitution (dS) rates to investigate signatures of selection on members of the NSP-like gene family compared with signatures on other larval-expressed orthologs. We also conducted comprehensive feeding experiments with 25 Brassicaceae plants to acquire patterns of host utilization in *Pieris* species to test if shifts in these patterns can be correlated with the evolution of NSP-like gene family members in *Pieris* butterflies. Additionally, we searched for functional gene groups with signatures of selection among the five *Pieris* species based on gene ontology (GO) and dN/dS analyses to identify potential genes related to host-plant detoxification which might be under positive or negative selection. By combining these approaches, we were able to investigate signatures of selection on ecologically important NSP-like gene family members and detoxification-related genes associated with host-plant utilization patterns in *Pieris* larvae (Fig. 2). Results provide important insights into the evolution of adaptive key innovations in *Pieris* butterflies.

**Fig. 2.**
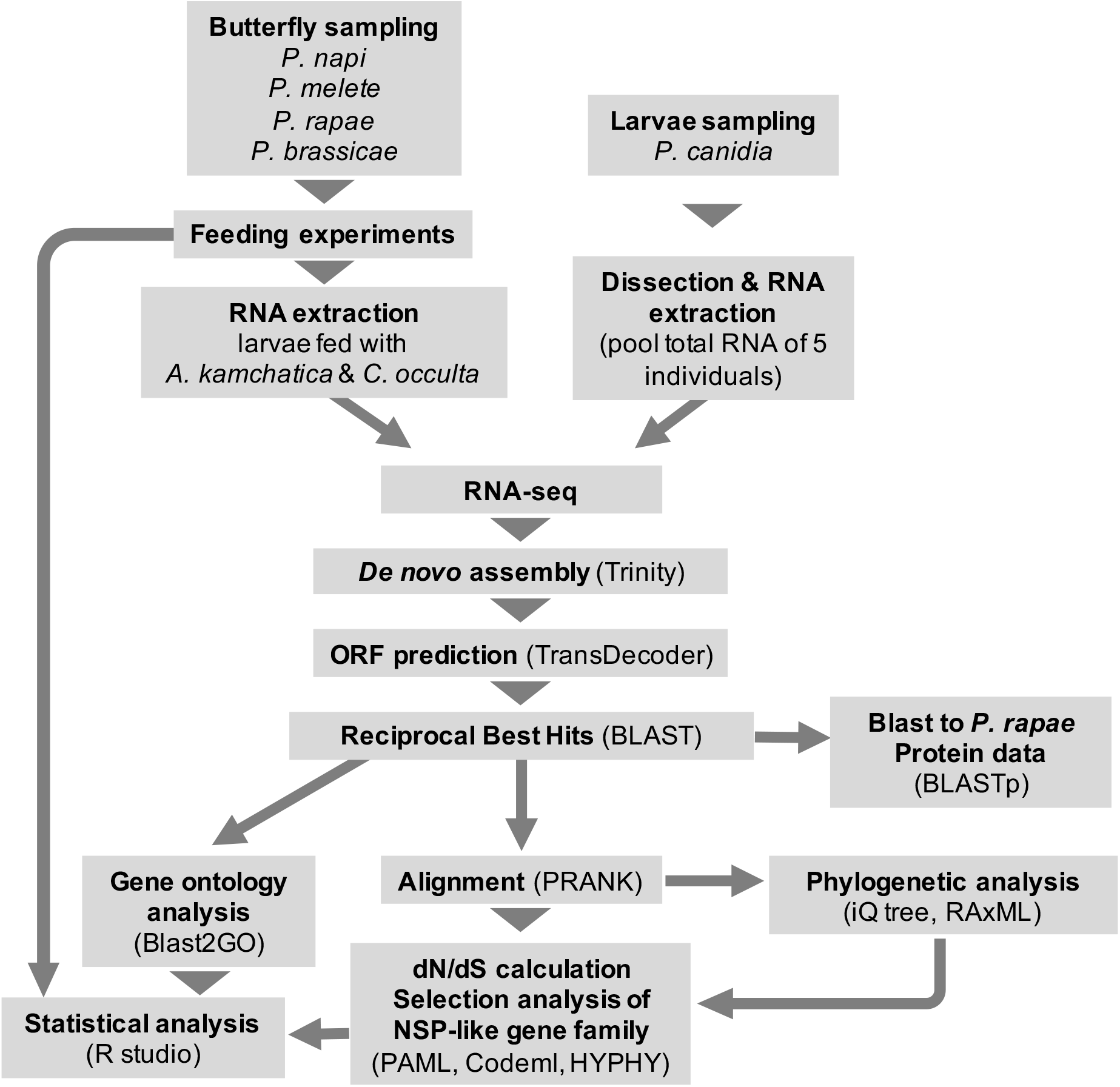
Analysis pipeline used to compare dN/dS ratios of NSP-like gene family members with all observed ortholog sets from the reciprocal best hit using BLAST across five *Pieris* butterflies. Signatures of selection on NSP-like gene family members were investigated in each phylogenetic branch and compared with the results of the feeding assay.

## Materials and Methods

### Feeding experiments

We used four *Pieris* butterfly species for the feeding assay, leaving out *P. canidia*. We collected 7– 10 female butterflies of three *Pieris* butterfly species (*P. napi*, *P. melete*, *P. rapae*) from wild populations in Chiba and Hokkaido, Japan. Most wild-caught female butterflies were already fertilized. We released the female butterflies into cages containing cabbage (*Brassica oleracea* var. *capitata*) or *Cardamine leucantha* under high-intensity light conditions, and waited for eggs to be laid. For *P. brassicae*, final-instar larvae were caught in the wild (Hokkaido, Japan), fed on cabbage and reared to the adult stage. After eclosion, 10 female butterflies were hand paired with males and eggs were collected as they were from the other species. Eggs of the four *Pieris* butterfly species were incubated at 25°C until they hatched.

For experimental plants, we collected seeds of 25 Brassicaceae plant species, covering a phylogenetically broad range (Table S1) (Beilstein, Al-Shehbaz, Mathews, & Kellogg, 2008; Couvreur et al., 2010; Franzke, Lysak, Al-Shehbaz, Koch, & Mummenhoff, 2011). We grew the plants in the greenhouse at 25°C, with 60% relative humidity and L16:D8. Plants were watered and fertilized every week with a 2000× diluted solution of Hyponex (N:P:K = 6:10:5; Hyponex, Osaka, Japan). After two months of cultivation, plants were used for the feeding experiments.

Neonate larvae were collected within 12 hours after they hatched for the feeding experiment. We transferred three neonate larvae for each plant using a soft-haired brush and replicated this twice for each plant species (*n* = 6). To minimize changes in the condition of the experimental plants, experimental trials were carried out within 5 days for all four *Pieris* species. We conducted feeding experiments under the same temperature and light conditions used for plant growth. We measured the weight of each larva individually (within 0.1 mg) after 120 hours of feeding and used the average weight of larval individuals from each plant species as an index of the performance of each *Pieris* butterfly species. We set the weight of dead larvae at 0.

Larval weights were standardized as z-scores to enable comparison between species. We calculated the mean scores of each plant treatment and used these for the comparative analysis. We conducted Pearson’s correlation test and hierarchical clustering analysis to assess differences in larval performances among the four *Pieris* species. The possible clustering was evaluated with the gap statistics (Tibshirani, Walther, & Hastie, 2001). All of these analyses were performed on R studio ver. 1.1.453 (RStudioTeam, 2016).

### RNA sequencing

From four *Pieris* butterfly species (*P. napi*, *P. melete*, *P. rapae, P. brassicae*), excluding *P. canidia*, we collected larvae that we used for the feeding experiments for transcriptome analysis (Figs. 1, 2). We used larvae that fed on *Arabidopsis kamchatica* and *Cardamine occulta* as representatives. The larvae were flash-frozen in liquid nitrogen and stored at −80 °C until RNA extraction. We selected a single representative larva for each of the four *Pieris* and plant species combinations, and RNA was extracted using the RNeasy Mini Kit (QIAGEN). RNA sample quantity and quality were checked by Agilent 2100 Bioanalyzer. Illumina libraries of individual larva were prepared by Sure Select Strand-Specific RNA Library Preparation Kit for Illumina Multiplexed Sequencing, and RNA sequencing was performed on an Illumina HiSeq 1500 Genome Analyzer platform using a 2 × 100bp paired-end approach. For *P. canidia*, we collected larvae directly from wild *Lepidium virginicum* on Yonaguni Island, Okinawa, Japan. The collected larvae were dissected, and gut tissues were stored at −80 °C in solution until RNA extraction. Five larvae were randomly selected, and RNA was extracted with the RNeasy Mini Kit (QIAGEN). *P. canidia* RNA concentrations were quantified on a Qubit 2 Fluorometer (Invitrogen), and a fraction of the RNA from each of the five larvae was pooled as a single sample for RNA-seq. Paired-end (2×150 bp) sequencing was performed by the Max Planck Genome Center Cologne on an Illumina HiSeq 2500 Genome Analyzer platform.

### *De novo* assembly, searching for reciprocal best hits (RBHs) using-BLAST

Acquired reads of RNA-seq data were pooled for each species after filtering out bad quality reads by trimmomatic with the following options (LEADING:10 TRAILING:10 SLIDINGWINDOW:4:20 MINLEN:40) (Bolger, Lohse, & Usadel, 2014). The quality of reads was checked by FastQC. Pooled reads were *de novo* assembled by Trinity ver. 2.0.6 (Grabherr et al., 2011). We used TransDecoder (http://transdecoder.github.io/) to predict open reading frames (ORFs) from the assembled contigs and subsequently looked for reciprocal best hits (RBHs) using BLAST alignment methods to analyze amino acid sequences (longer than 100 amino acids) predicted by TransDecoder (Camacho et al., 2009; Cock, Chilton, Grüning, Johnson, & Soranzo, 2015). We used RBH BLAST software with default settings on all possible species pairs (10 pairs) and subsequently extracted *P. rapae* orthologs from this RBH result and ran blastp on the amino acid sequences against a *P. rapae* protein database to confirm the ORF prediction from TransDecoder. Orthologs in the RBH result without any BLAST hits to the *P. rapae* protein database were removed since these amino acid sequences may have resulted from wrong ORF predictions by TransDecoder. We used PRANK to conduct codon-based alignment of each ortholog set acquired from the RBH result (Loytynoja & Goldman, 2005).

### Phylogenetic tree construction

We reconstructed a phylogeny of the five *Pieris* species using the transcriptome data by concatenating all aligned ortholog nucleotide sequences into one sequence for each species, generating an Maximum Likelihood (ML) phylogenetic tree by IQ-tree (Nguyen, Schmidt, Von Haeseler, & Minh, 2015) after removing gaps with TrimAl (2063074 bp remaining) (Capella-Gutiérrez, Silla-Martínez, & Gabaldón, 2009). We used the GTR + gamma substitution model and set ultrafast bootstrap approximation iterations as 1000, using -bnni options to construct a phylogeny of the five *Pieris* species (Hoang, Chernomor, Von Haeseler, Minh, & Vinh, 2018).

### Analysis of NSP-like gene family members

We used each aligned orthologous gene for calculating species pairwise dN/dS ratios using PAML 4.8 (Yang, 2007). We used runmode = −2 and NSsites = 0 option in codeml from PAML and calculated pairwise dN/dS ratios based on the Nei & Gojobori method (Nei & Gojobori, 1986). We averaged acquired species pairwise dN/dS ratios for each ortholog to infer putative positive selection among the genus. The dN/dS values of NSP-like gene family members were compared with the entire dN/dS distributions of all ortholog sets in species pairwise associations and also in an averaged dN/dS scale among *Pieris*.

We used the branch-site model to identify cases of positive selection on NSP-like gene family members in a specific branch. We prepared molecular phylogeny of NSP gene family members by RAxML (Stamatakis, 2014) and tested all branches using codeml model 2 with NSsites = 2 option and ran an alternative model; varied dN/dS ratios across sites as well as lineages were allowed (fixed_omega = 0), and null model; fixed dN/dS (fixed_omega = 1). We conducted a likelihood ratio test (LRT) with the chi-square distribution to evaluate significant differences between the alternative and null models. We also used adaptive Branch-Site Random Effects Likelihood (aBSREL) analysis for the NSP-like gene members among the five *Pieris* species and tested all branches to identify the signatures of positive selection (Smith et al., 2015). Acquired *P* values were corrected with false discovery rates (FDRs) in each analysis. Signs of positive selection on each site were identified by the Bayes empirical Bayes (BEB) analysis (0.90 cut offs). The aBSREL analyses were performed in HyPhy implemented in the datamonkey web server (Kosakovsky Pond, Frost, & Muse, 2005; Weaver et al., 2018).

### GO annotation and evolutionary tests

We used *P. rapae* contigs from the RBH result for GO annotation and ran these genes against the NCBI non-redundant protein sequence database in Galaxy (Blastx, e-value = 10e-4). We subsequently used the Blast2GO platform to load the resulting Blast-xml file and to conduct mapping and annotation steps based on the BLAST result for acquiring GO annotations for each contig (Götz et al., 2008). To test significantly elevated or decreased dN/dS ratios among genes associated with specific GO terms, we selected those that contained at least 20 orthologs and tested their dN/dS distributions with those of all the observed orthologs (background) using a Wilcoxon test. All statistical analyses were performed in R studio ver. 1.1.453 and *P* values acquired were adjusted by FDR (RStudioTeam, 2016).

## Results

### Performance of four *Pieris* butterflies on 25 Brassicaceae plants

We obtained larval weights for four *Pieris* butterfly species (*Pieris napi*, *P. melete*, *P. rapae* and *P. brassicae*) feeding on 25 different Brassicaceae plant species. Analysis showed that the larval performances of the four *Pieris* species could be clustered into two groups: the *P. napi-P. melete* group and the *P. rapae-P. brassicae* group. The gap statistics for the given number of clusters were as follows: *Gap*_1_ =0.080, *Gap*_2_ = 0.135, *Gap*_3_ = 0.119, *Gap*_4_ = 0.123 (Fig. 3). For instance, we observed that *P. napi* and *P. melete* performed better on *Arabis hirsuta* or *Turritis glabra*, whereas *P. rapae* and *P. brassicae* did better on *Thlaspi arvense* than the other two species (Fig. 3). In addition to this trend, the larvae of four *Pieris* butterfly species also performed similarly. We observed that all four *Pieris* species performed better on *Cardamine occulta* than on the other plant species tested and did not perform well on *Erysimum cheiranthoides* or *Berteroa incana* (Fig. 3).

**Fig. 3.**
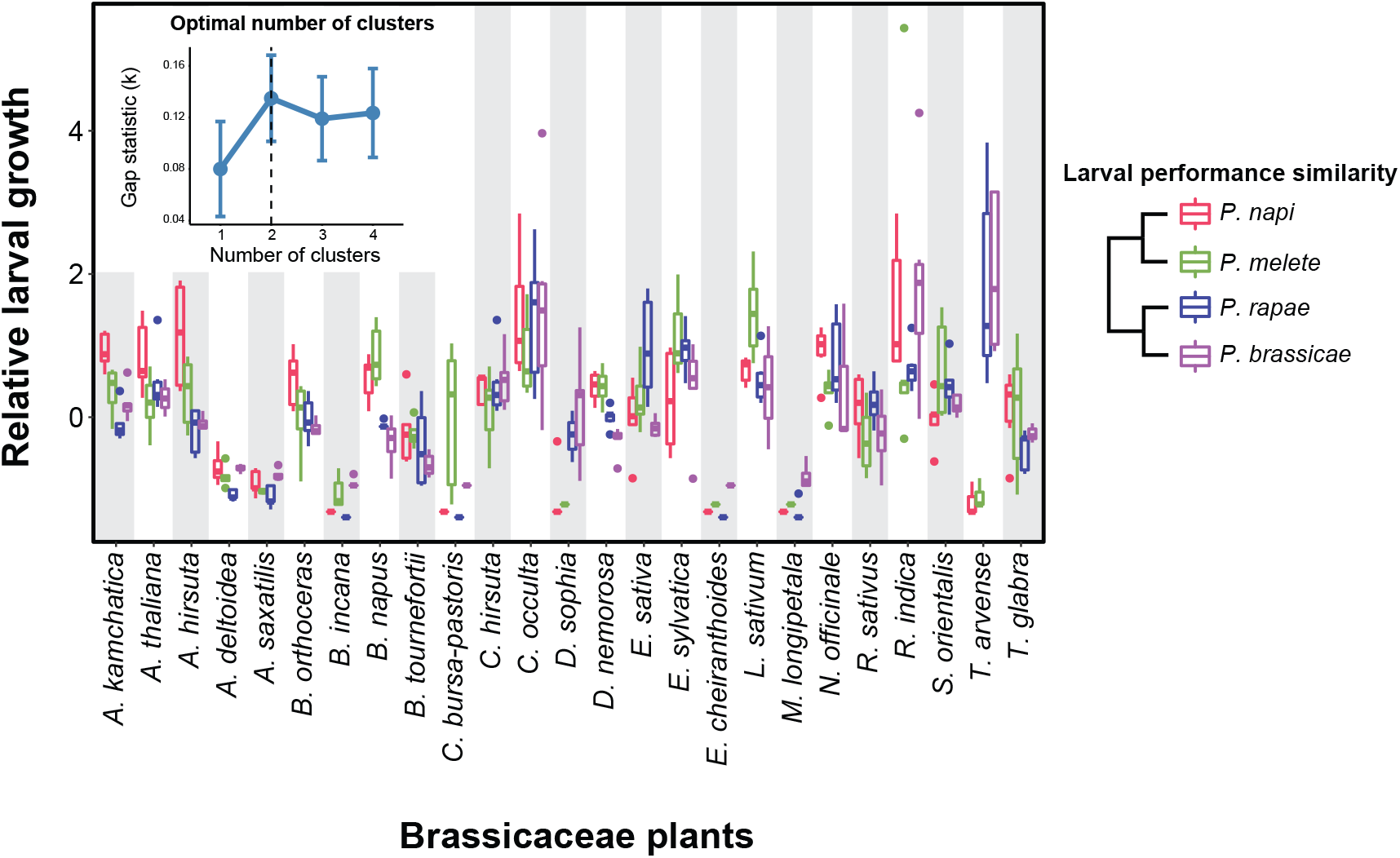
Feeding assays of four *Pieris* butterfly larvae on 25 different Brassicaceae plants (*n* = 6). The four *Pieris* butterfly species generally grew better on *Cardamine occulta* but could not use *B. incana* or *E. cheiranthoides* as optimal hosts. Overall larval performance patterns of the four *Pieris* species could be clustered in two groups: *P. napi – melete* and *P. rapae – brassicae*. *P. napi* and *P. melete* larvae grew better on *Arabis hirsuta* or *Turritis glabra*, and *P. rapae* and *P. brassicae* larvae grew better on *Thlaspi arvense*.

### RNA-seq, reciprocal best hit (RBH) BLAST analysis of *Pieris* butterflies

We obtained 32-40 million Illumina 100 bp pair-end reads for the four species (*P. napi*, *P. melete*, *P. rapae* and *P. brassicae*) and 64 million Illumina 150 bp pair-end reads for *P. canidia*. *De novo* transcriptome assemblies using Trinity resulted in 64,279; 62,054; 59,327; 53,004; and 149,481 contigs, and in N50 values of 2,048 bp; 2,132 bp; 2,060 bp; 2,594; and 2,075 bp for *P. napi*, *P. melete*, *P. rapae P. brassicae*, and *P. canidia* respectively. Using RBH BLAST on the five *Pieris* species, we obtained transcriptome data resulted in 2723 ortholog sets.

### Identifying signatures of selection on NSP-like gene family members

We calculated dN/dS ratios for all ortholog sets in the 10 *Pieris* species pairs used with PAML 4.8 (Yang, 2007) and averaged these values. The complete distribution of averaged dN/dS values is shown in Figure 4 (mean dN/dS = 0.10486). The averaged dN/dS values of NSP-like gene family members are as follows: dN/dS_NSP_ = 0.324, dN/dS_MA_ = 0.188 and dN/dS_SDMA_ = 0.125. The dN/dS value of NSP is located in the top 2.72% of the entire dN/dS distribution, whereas MA and SDMA values are lower (MA 11.4%, SDMA 23.4%). We also found a similar pattern between species, where NSPs were in the top 5% in 5 pairs out of 10 (*napi – rapae, napi – canidia, melete – rapae, melete – canidia, rapae – canidia*) and in the top 5.5% in two pairs (*napi – brassicae, melete – brassicae*); MAs and SDMAs were not ranked in the top 5 % (Fig. 5). In most cases, NSPs had the highest value, MAs had higher dN/dS value compared to SDMAs, and the order of dN/dS values of NSP-like gene family members was NSP > MA > SDMA in 8 out of 10 species pairs (Fig. 5).

**Fig. 4.**
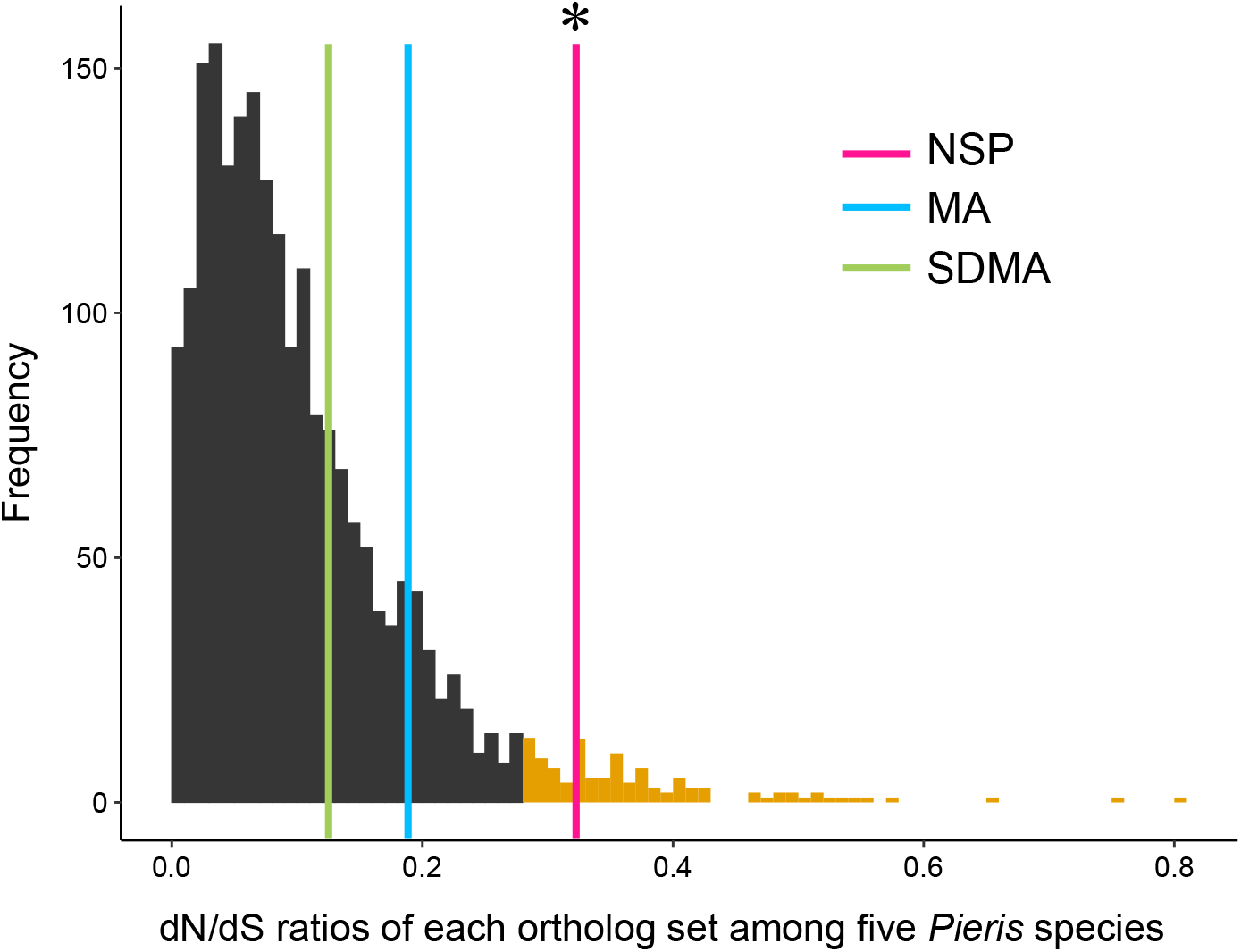
The averaged distribution of dN/dS values within each ortholog among the five *Pieris* species (*n* = 2723). The top 5% values in the histogram are colored orange. The vertical lines show dN/dS values of NSP-like gene family members; NSP (pink) = 0.324, MA (blue) = 0.188 and SDMA (green) = 0.125. ‘*’ shows the line is in the top 5%. The dN/dS values of NSP are located in the top 5% of the entire dN/dS distribution (NSPs are located in 2.72%), whereas those of MAs and SDMAs are not (MAs 11.4%, SDMAs 23.4%).

**Fig. 5.**
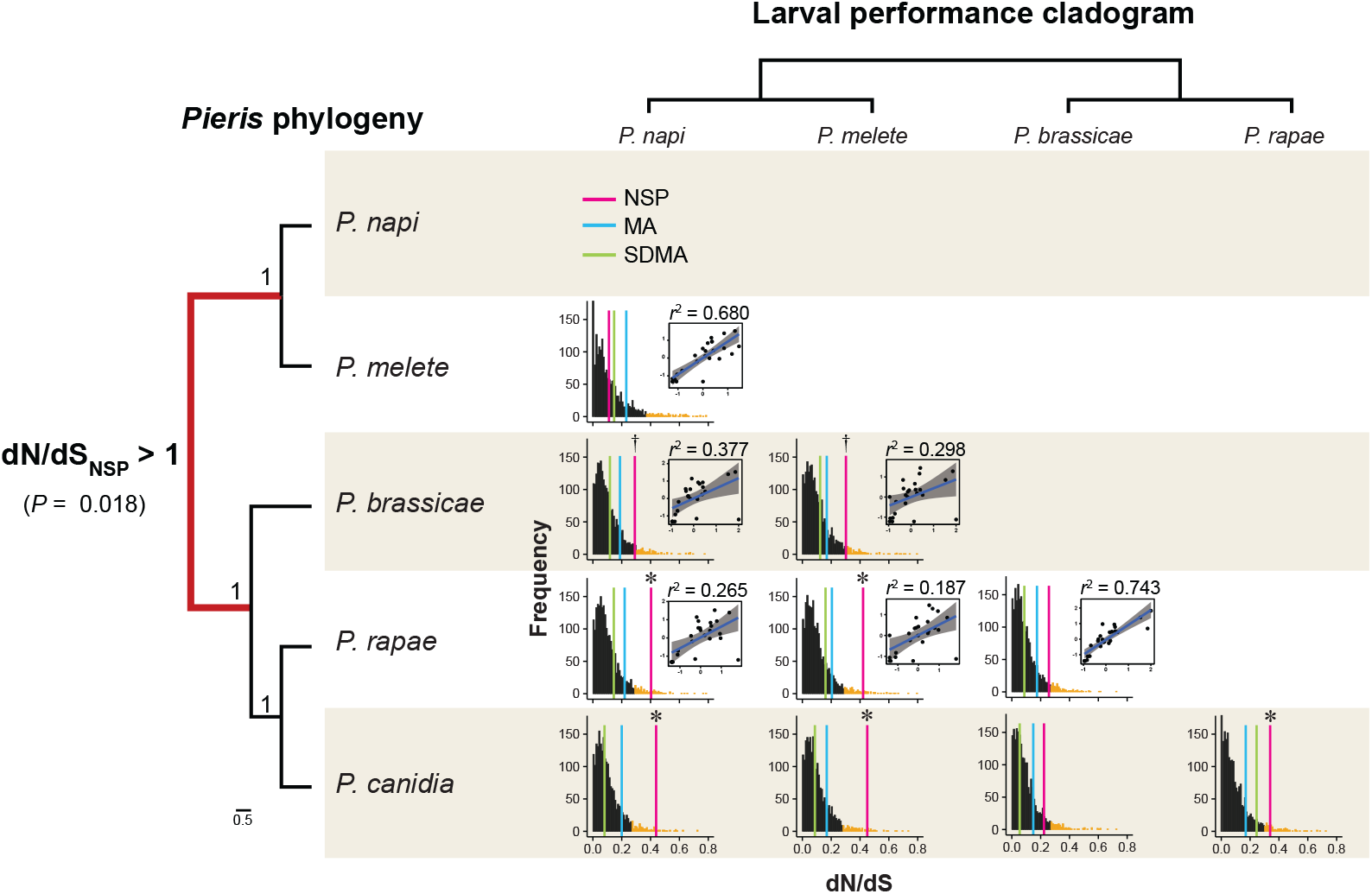
The species pairwise dN/dS values compared with the results of feeding behavior of *Pieris* butterflies. The histograms showed distributions of species pairwise dN/dS values of entire ortholog sets (showing only from 0 ≤ dN/dS ≤ 0.8 for displaying in scale and the top 5% are colored orange). The positions of each NSP-like gene family member are highlighted with colored vertical lines (NSP: pink, MA: blue, SDMA: green). Symbols on the lines show ranking (‘*’: in the top 5%, ‘†’: in the top 5.5%). The scatterplots show the larval growth of each pair of *Pieris* species except for *P. canidia*. A phylogenetic tree was reconstructed by all the aligned contigs of the five *Pieris* species with bootstrapping support in each node. We found a signature of positive selection in NSPs at the *P. melete* and *P. napi* branch (left phylogeny: *P* = 0.018, LRT), which is also supported by significantly elevated dN/dS values of NSP in the species pairs at the phylogenetic branch (*P. napi -. P. rapae, P. napi - P. brassicae, P. melete - P. rapae, P. melete - P. brassicae*). Furthermore, dissimilarities in larval performance correlated with elevated dN/dS values of NSPs.

### Signature of clade-specific positive selections on NSP associated with larval performance

We reconstructed Japanese *Pieris* phylogeny using the transcriptome data by concatenating all RBH ortholog sets. The obtained results showed a highly supported *P. napi*–*P. melete* clade, and *P. rapae*–*P. canidia* clade (Fig. 5). We performed the branch-site model approach by codeml and found a signal of positive selection on NSPs at the *P. napi*–*P. melete* branch (FDR adjusted *P* = 0.0178, LRT), however, we found no sign of positive selection at other branches or in MA or SDMA genes (Fig. 5, Table 1). The BEB analysis suggested that two codon sites had signs of positive selection in NSPs in this branch (Table 1, posterior probability > 0.9). These sites were located in second and third domains of NSPs (position 421 and 503 in the amino acid sequence) and close to the positively selected sites identified in previous work (positions 379 and 523) (Heidel-Fischer et al., 2010). The aBSREL analyses also detected a signature of positive selection on NSP genes only at the *P. napi*–*P. melete* branch (FDR adjusted *P* = 0.010), whereas no branch-specific positive selection was detected in MA and SDMA genes (Table 2).

**Table 1.**
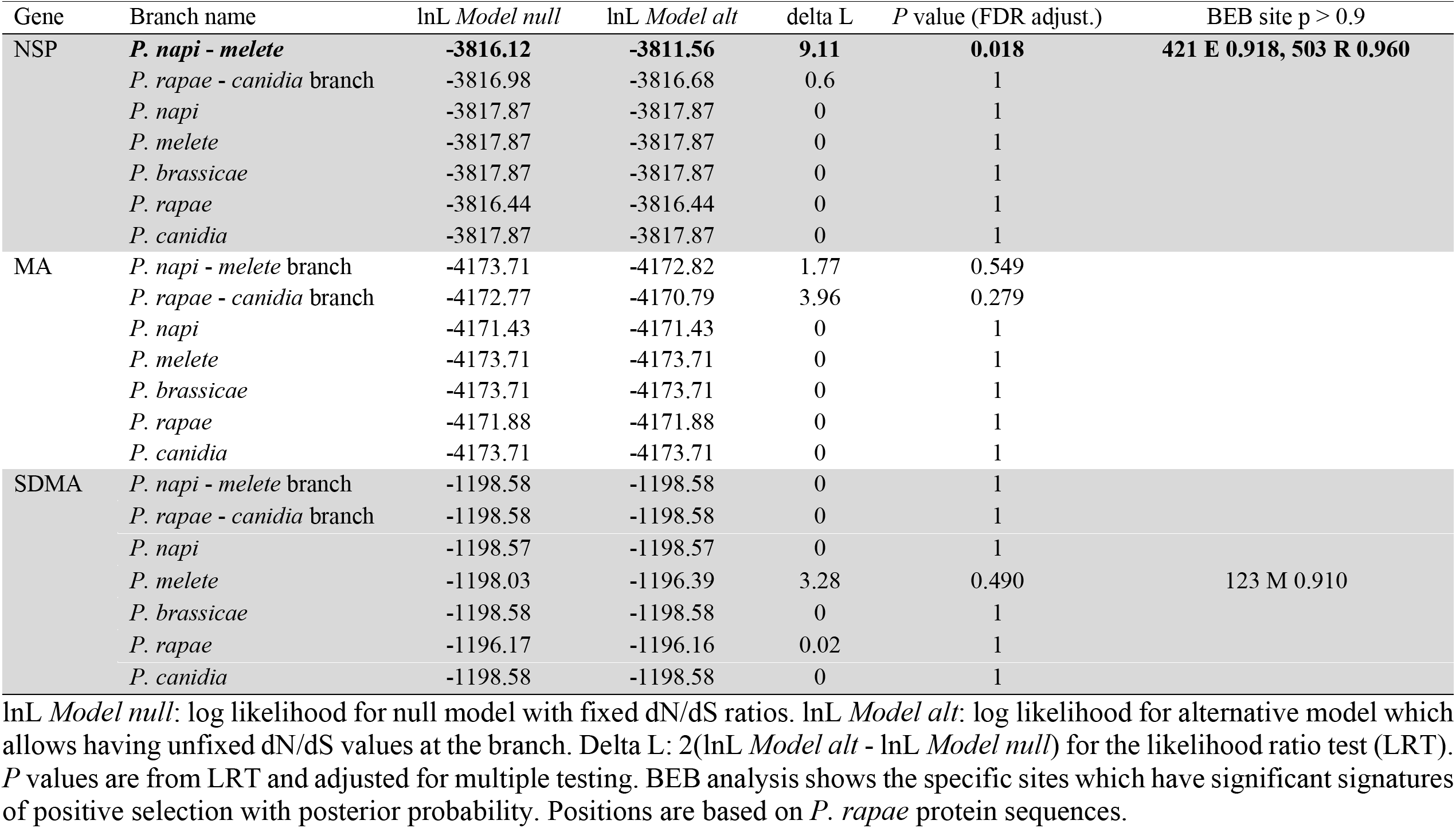
Branch-specific selection tests on NSP-like gene family by codeml.

**Table 2.** Branch-specific selection tests on NSP-like gene families by a BSREL.

| Gene | Branch name | LRT | *P* value |
| --- | --- | --- | --- |
| NSP | ***P. napi* - *melete* branch** | **10.96** | **0.010** |
|  | *P. rapae* - *canidia* branch | 5.06 | 0.067 |
|  | *P. napi* | 0 | 1 |
|  | *P. melete* | 5.46 | 0.067 |
|  | *P. brassicae* | 2.33 | 0.208 |
|  | *P. rapae* | 0 | 1 |
|  | *P. canidia* | 0 | 1 |
| MA | *P. napi* - *melete* branch | 3.02 | 0.967 |
|  | *P. rapae* - *canidia* branch | 4.43 | 0.967 |
|  | *P. napi* | 0 | 1 |
|  | *P. melete* | 0 | 1 |
|  | *P. brassicae* | 0.96 | 1 |
|  | *P. rapae* | 0 | 1 |
|  | *P. canidia* | 4.27 | 0.967 |
| SDMA | *P. napi* - *melete* branch | 0 | 1 |
|  | *P. rapae* - *canidia* branch | 0.49 | 1 |
|  | *P. napi* | 0 | 1 |
|  | *P. melete* | 4.38 | 1 |
|  | *P. brassicae* | 0 | 1 |
|  | *P. rapae* | 0 | 1 |
|  | *P. canidia* | 0 | 1 |

LRT: Likelihood ratio test statistic for selection. *P* values are adjusted by false discovery rates.

### GO terms with elevated dN/dS ratios among five *Pieris* butterflies

After GO annotations of all *P. rapae* RBH contigs, we obtained 1457 GO terms in our datasets. These included 680 related to biological process, 540 to molecular function, and 237 to cellular component GO terms. We conducted the Wilcoxon test for the GO terms, which have more than 20 assigned orthologs, and the result revealed that one biological process --“proteolysis” -- and two processes associated with molecular function --“hydrolase activity” and “serine-type endopeptidase activity” --had significantly elevated dN/dS values when compared to the entire dN/dS distribution of all contigs (Fig. 6, Table 3). This test also showed that 13 GO terms had significantly lower dN/dS values in the three categories (Table 3). These lower dN/dS GO terms included “regulation of transcription, DNA-templated,” “ribosome biogenesis,” and “translation” in biological process; “ATP binding,” “structural constituent of ribosome,” “GTP binding,” “calcium ion binding,” “DNA-binding transcription factor activity” and “sequence-specific DNA binding” in molecular functions; and “nucleus,” “cytoplasm,” “ribosome” and “transcription factor complex” in the category of GO cellular components.

**Fig. 6.**
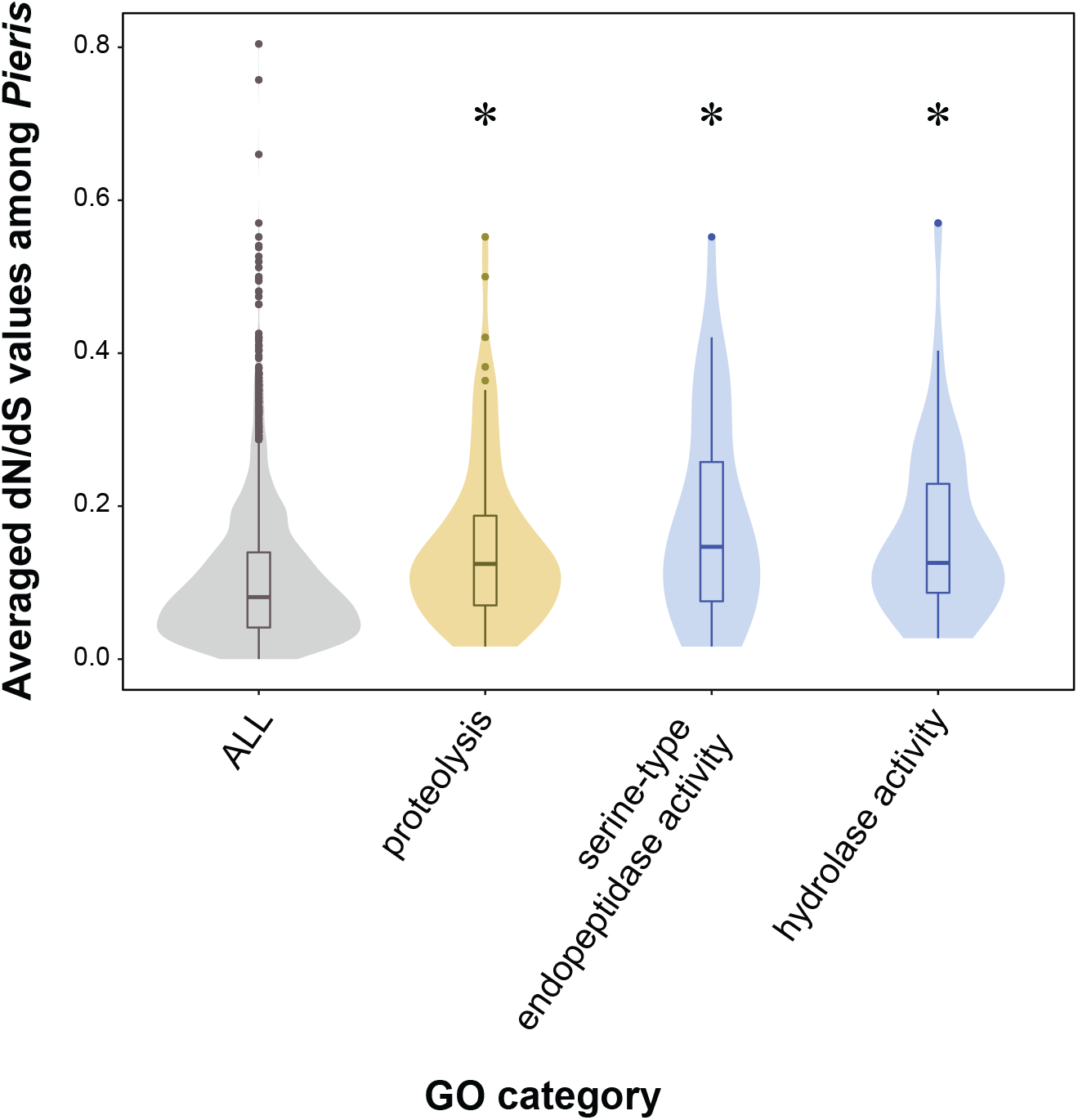
GO terms which have significantly elevated dN/dS values compared to those of entire ortholog sets among *Pieris*. Elevated dN/dS values were observed in “proteolysis” from biological process (orange), and “serine-type endopeptidase activity” and “hydrolase activity” from molecular function (blue) compared to the entire distribution of all the observed contigs among *Pieris* (gray). Comparisons with other enriched GO terms are shown in Table 3. ‘*’: *P* values are adjusted by FDR; *P* ≤ 0.05, Wilcoxon test.

**Table 3.**
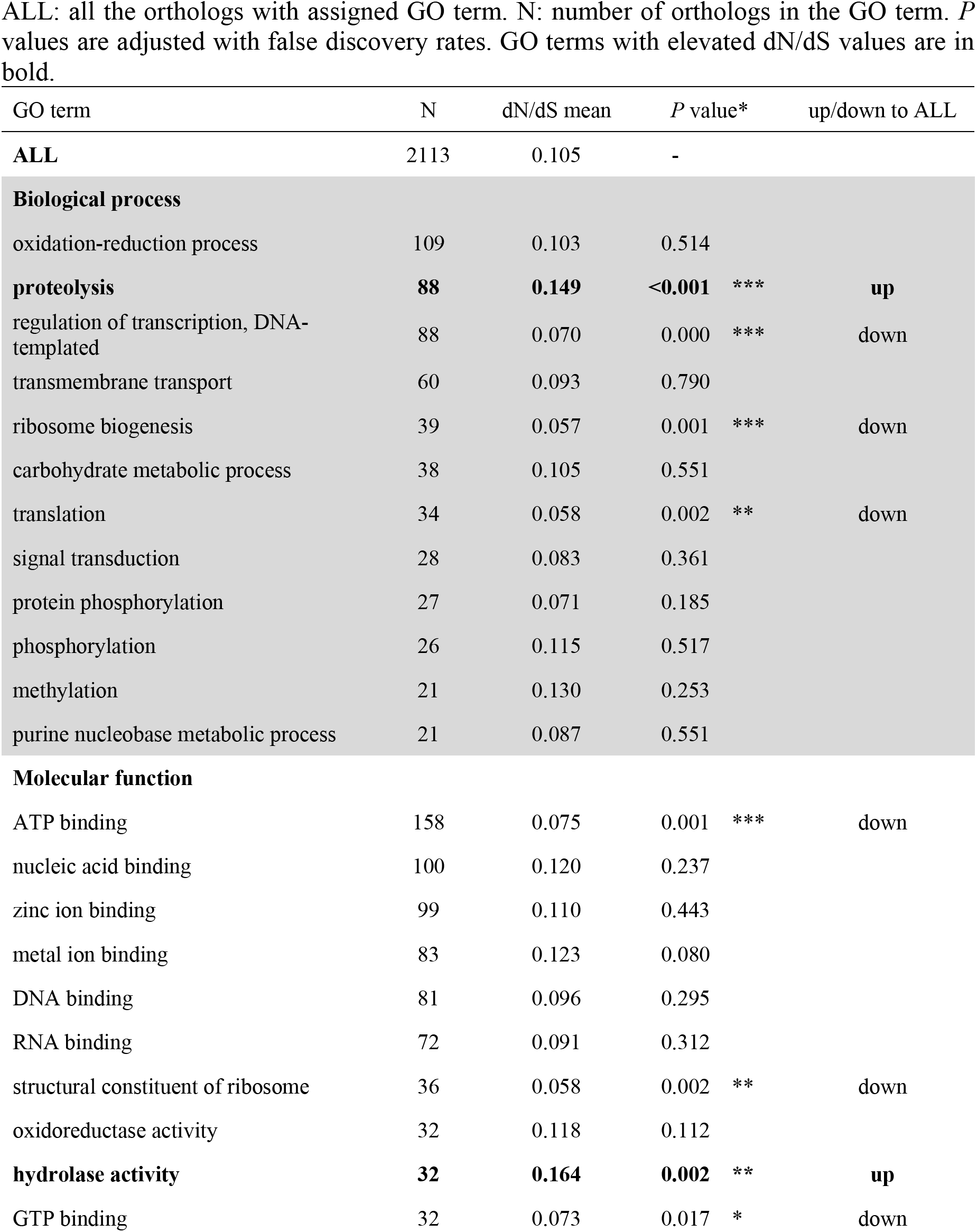
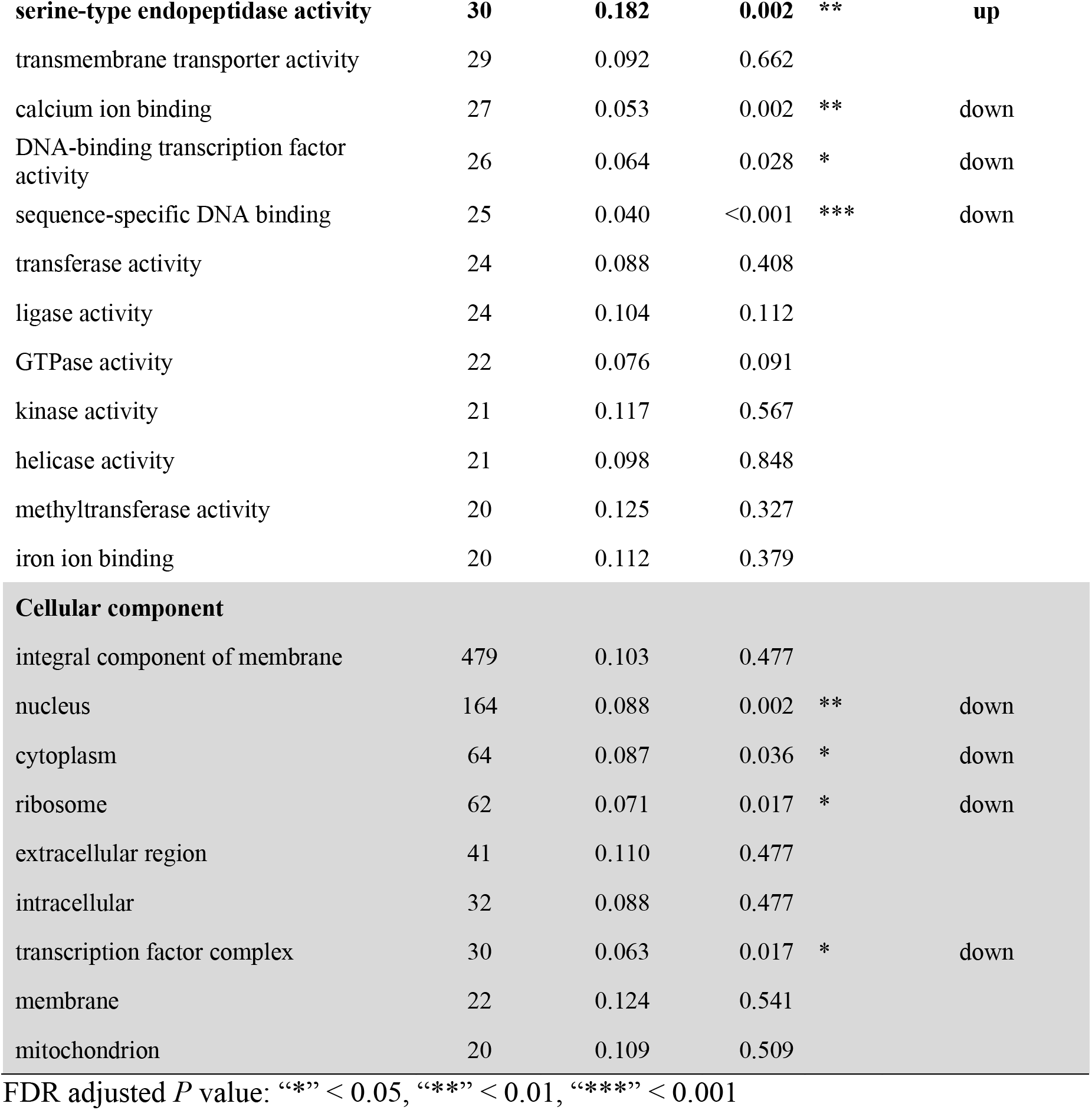
GO terms with elevated or decreased dN/dS values corresponding to entire orthologs. ALL: all the orthologs with assigned GO term. N: number of orthologs in the GO term. *P* values are adjusted with false discovery rates. GO terms with elevated dN/dS values are in bold.

| GO term | N | dN/dS mean | *P* value* | up/down to ALL |
| --- | --- | --- | --- | --- |
| **ALL** | 2113 | 0.105 | - |  |
| **Biological process** |  |  |  |  |
| oxidation-reduction process | 109 | 0.103 | 0.514 |  |
| **proteolysis** | **88** | **0.149** | **<0.001** *** | **up** |
| regulation of transcription, DNA-templated | 88 | 0.070 | 0.000 *** | down |
| transmembrane transport | 60 | 0.093 | 0.790 |  |
| ribosome biogenesis | 39 | 0.057 | 0.001 *** | down |
| carbohydrate metabolic process | 38 | 0.105 | 0.551 |  |
| translation | 34 | 0.058 | 0.002 ** | down |
| signal transduction | 28 | 0.083 | 0.361 |  |
| protein phosphorylation | 27 | 0.071 | 0.185 |  |
| phosphorylation | 26 | 0.115 | 0.517 |  |
| methylation | 21 | 0.130 | 0.253 |  |
| purine nucleobase metabolic process | 21 | 0.087 | 0.551 |  |
| **Molecular function** |  |  |  |  |
| ATP binding | 158 | 0.075 | 0.001 *** | down |
| nucleic acid binding | 100 | 0.120 | 0.237 |  |
| zinc ion binding | 99 | 0.110 | 0.443 |  |
| metal ion binding | 83 | 0.123 | 0.080 |  |
| DNA binding | 81 | 0.096 | 0.295 |  |
| RNA binding | 72 | 0.091 | 0.312 |  |
| structural constituent of ribosome | 36 | 0.058 | 0.002 ** | down |
| oxidoreductase activity | 32 | 0.118 | 0.112 |  |
| **hydrolase activity** | **32** | **0.164** | **0.002** ** | **up** |
| GTP binding | 32 | 0.073 | 0.017 * | down |

|  |  |  |  |  |  |
| --- | --- | --- | --- | --- | --- |
| **serine-type endopeptidase activity** | **30** | **0.182** | **0.002** | **\*\*** | **up** |
| transmembrane transporter activity | 29 | 0.092 | 0.662 |  |  |
| calcium ion binding | 27 | 0.053 | 0.002 | ** | down |
| DNA-binding transcription factor activity | 26 | 0.064 | 0.028 | * | down |
| sequence-specific DNA binding | 25 | 0.040 | <0.001 | *** | down |
| transferase activity | 24 | 0.088 | 0.408 |  |  |
| ligase activity | 24 | 0.104 | 0.112 |  |  |
| GTPase activity | 22 | 0.076 | 0.091 |  |  |
| kinase activity | 21 | 0.117 | 0.567 |  |  |
| helicase activity | 21 | 0.098 | 0.848 |  |  |
| methyltransferase activity | 20 | 0.125 | 0.327 |  |  |
| iron ion binding | 20 | 0.112 | 0.379 |  |  |
| **Cellular component** |  |  |  |  |  |
| integral component of membrane | 479 | 0.103 | 0.477 |  |  |
| nucleus | 164 | 0.088 | 0.002 | ** | down |
| cytoplasm | 64 | 0.087 | 0.036 | * | down |
| ribosome | 62 | 0.071 | 0.017 | * | down |
| extracellular region | 41 | 0.110 | 0.477 |  |  |
| intracellular | 32 | 0.088 | 0.477 |  |  |
| transcription factor complex | 30 | 0.063 | 0.017 | * | down |
| membrane | 22 | 0.124 | 0.541 |  |  |
| mitochondrion | 20 | 0.109 | 0.509 |  |  |

FDR adjusted *P* value: “\*” < 0.05, “\*\*” < 0.01, “\*\*\*” < 0.001

## Discussions

Focusing on five Japanese *Pieris* butterflies, we tested host-plant performance and investigated signatures of selection on NSP-like genes, which are a key innovation of these butterflies to overcome the GLS defense system of their Brassicales host plants (Edger et al., 2015; Wheat et al., 2007). We acquired RBH ortholog sets expressed in larvae of the five *Pieris* species based on transcriptome data and compared the calculated dN/dS ratios of each ortholog in order to investigate the effect of evolutionary forces on NSP-like gene family members. We also combined ecological approaches for acquiring performance data on larvae of *Pieris* species by conducting comprehensive feeding experiment using 25 Brassicaceae plant species. These approaches yielded four major findings. First, we observed that *Pieris* species showed clade-specific differences in larval host performance. Second, we observed that NSP genes had significantly elevated dN/dS ratios compared to other genes in the five *Pieris* species, including members of the same gene family, MAs and SDMAs. Third, evidence of positive selection on NSPs was observed at a phylogenetic branch which showed differences in larval performance according to our feeding assays. Last, we observed significantly elevated dN/dS ratios in GO terms which are associated with potential detoxification-related genes in *Pieris* larvae.

According to our feeding experiments with four Japanese *Pieris* species (*P. napi*, *melete*, *rapae* and *brassicae*) and 25 Brassicaceae plant species, *P. napi* and *P. melete* larvae performed similarly, as did *P. rapae* and *P. brassicae* larvae (Fig. 3). Observations in the field suggest that these four *Pieris* species have slightly different host preferences: *P. napi* and *P. melete* feed on wild and montane Brassicaceae plants, such as *Arabis* or *Turritis*, and *P. rapae* and *brassicae* use Brassicaceae crops more often than the other two species (Fig. 1) (Harvey, Poelman, & Gols, 2010; Ohsaki & Sato, 1994). Thus, our results confirm the field observations (Fig. 3). Phylogenetic analysis showed that the *P. napi* and *P. melete* clade was strongly supported, and the species phylogeny seemed to correspond with larval performance (Fig. 5), suggesting that the larval host preferences of the four *Pieris* butterflies are phylogenetically conserved. In this study, we did not perform any physical or chemical defense analyses on the different Brassicaceae plants species we used; however, a number of previous studies revealed that the GLS profiles of Brassicaceae plants can differ dramatically among Brassicaceae species (Agerbirk & Olsen, 2012; Fahey, Zalcmann, & Talalay, 2001; Olsen et al., 2016). Our results suggest that *Pieris* species might not always be capable of fully adapting to the defenses in the ranges of their potential host plants and so likely evolved to feed on a subset of Brassicaceae plants.

Comparing averaged dN/dS ratios among all species pairs for each ortholog, we found that only NSPs had higher dN/dS values among NSP-like gene family members (Fig. 4). Although we filtered out a number of genes by RBH processes and therefore compared only a subset of the entire orthologs, our finding strongly suggests that NSPs are under positive selection – or, more relaxed purifying selection -- among the five *Pieris* butterfly species. In our interspecies dN/dS comparison, we also observed that NSPs had higher dN/dS values than the other ortholog sets in most of the species pairs (they were located in the top 5.5% in 7 out of 10 species pairs), supporting the hypothesis of positive selection on NSPs in this genus (Fig. 5). In previous research, which calculated dN/dS values from partial NSP sequences of *P. rapae* and *P. brassicae* with 70 other genes, higher dN/dS values of NSPs were observed (dN/dS = 0.25 ranked in the top 5%) (Heidel-Fischer et al., 2010). We found that dN/dS values in our dataset from entire NSP mRNA sequences of this species pair were 0.257, which ranked in the top 6.06 %, thus supporting previous findings. Interestingly, we also found that in most cases MAs had lower dN/dS values compared to NSPs (in both averaged dN/dS ratios and interspecies comparisons) (Figs. 4, 5), and their dN/dS values did not reach the top 5%, suggesting that in this genus, MAs are under stronger purifying selection than are NSPs. NSPs and MAs are known as paralogs, and only NSP was confirmed to have GLS-disarming activity in *Pieris*. However, MAs also disarm GLSs in another Brassicaceae-feeding Pierid genus, *Anthocharis*, which has only MAs (Edger et al., 2015); this overlap strongly suggests that in *Pieris* MAs act like NSPs. Our results show that selection on these two paralogous genes, both of which have similar structure and can potentially disarm GLSs, can differ strikingly. This could imply that these paralogs have differentially functionalized in *Pieris*, where NSPs have more derived functions, whereas MAs have more conservative functions. DN/dS values of SDMAs were lowest among all NSP-like gene family members and also had similar values compared to the average of all the orthologous sets. This similarity suggests that SDMAs are under strong purifying selection and have a conserved function in *Pieris*. Expressed in the gut, SDMAs are known to be found in all Lepidoptera, supporting the hypothesis that their function is related to digestion (Fischer et al., 2008; Randall et al., 2013).

Using the branch-site model analysis by codeml, we detected evidence of positive selection only in NSPs at the *P. napi*-*melete* branch (Table 1). Testing all possible branches of all NSP-like gene family members with aBSREL, we detected signatures of positive selection in NSPs only at this branch (Table 2). Based on our comprehensive feeding experiment and phylogenetic analyses, we found that the *P. napi*-*melete* branch had different host preferences from *P. rapae* and *P. brassicae* (Figs. 2, 4). These results suggest that host-plant preferences in *Pieris* were associated with the evolution of NSPs but not MAs or SDMAs. In this study, we did not test the functional differences of NSPs among the five *Pieris* species. Furthermore, we could not determine whether the differences in larval performances that we observed among the four *Pieris* species were caused by the dissimilarity among the GLS profiles of the host plants. However, our findings imply a strong relationship between the evolution of NSPs and host-utilization patterns among *Pieris* butterflies. Moreover, it is also interesting that only NSPs showed this signature of selection, suggesting that NSPs have been functionalized to detoxify GLSs specific to certain plant species; in contrast, MAs may have evolved to disarm the widespread types of GLSs such as are found universally across Pieridae host plants. In addition, we found positively selected sites in the second and third domains of NSPs, and in earlier population genetic work using *P. rapae* (Heidel-Fischer et al., 2010). Although the molecular mechanisms of the GLS-disarming function of NSPs and MAs are still unclear, our results suggest that the second and third domains of NSPs are important for substrate specificity.

Besides individual NSP gene family members, elevated dN/dS values were also more broadly observed among the five *Pieris* butterflies in several GO categories, including “proteolysis” (biological process), and “serine-type endopeptidase activity” and “hydrolase activity” (molecular function). In Lepidopteran larvae, most of the digestive enzymes are involved in proteolysis (Simon et al., 2015) and several classes of digestive enzymes are necessary for insect herbivores to acquire essential nutrients in appropriate amounts (Broadway, 1989). In *Pieris*, these proteolytic activities were dominated by serine endopeptidases (Broadway, 1996). Since plants also have varied species-specific protease inhibitors to inhibit protease activity in herbivores, herbivores need to have evolved inhibitor-resistant proteinases as a counter adaptation (Bolter & Jongsma, 1997). Our findings showed signs of positive selection in protease-related genes among five *Pieris* species, suggesting that these genes have accumulated more functional changes as a consequence of interactions with plants in their specific host-plant ranges. A number of genes with hydrolase activity are included in genes related to detoxification in herbivores (Simon et al., 2015). Previous research has uncovered differential gene regulation of this GO term member in several herbivore species responding to different host-plants (Schweizer, Heidel-Fischer, Vogel, & Reymond, 2017). Therefore, a sign of positive selection or relaxed purifying selection on this GO member may also be associated with the host-plant spectra in *Pieris* butterflies.

To uncover the co-evolutionary diversification of plants and herbivores, it is important to understand the molecular interactions between all involved partners. We found the signature of positive selection on NSPs in a Pieridae genus, *Pieris*, associated with respective host-plant usage. It seems that the evolution of host-plant adaptive genes is correlated with patterns of host-plant usage in this *Pieris* butterfly genus. Moreover, we also observed that MAs, which are paralogs of NSPs, are under more strict purifying selection than NSPs. Our findings combine results from genetic and ecological assays to focus on how the evolution of these two paralogous genes may affect the arms-race between Brassicales and *Pieris* butterflies and their consequent diversification. Functional assays focusing on selected sites will increase our understanding of the evolution and functional differentiation of NSPs and MAs and how *Pieris* adapted evolutionarily to diverse glucosinolates in their host plants.

## Acknowledgements

We are grateful to Takashi Tsuchimatsu for useful discussions and comments on this study. We thank Emily Wheeler, Boston, for editorial assistance. This work was supported by a Grant-in-Aid for Scientific Research from the Japan Society for the Promotion of Science (15J00320 to Y.O.) and partially by Max-Planck-Gesellschaft.

## Data Accessibility

The RNA-seq short read data have been deposited in the EBI short read archive (SRA) with the following sample accession numbers: ERX2829492-ERX2829499. The complete study can also be accessed directly using the following URL: http://www.ebi.ac.uk/ena/data/view/PRJEB29048.

## Author Contributions

Y.O., A.S., and N.T. carried out the laboratory work. Y.O., M.M., H.H.F. and H.V. conceived, designed and coordinated the study. Y.O., M.M., H.H.F. and H.V. wrote the manuscript. All authors, drafted parts of the manuscript, gave approval for publication and agree to be accountable for the content.

## References

Agerbirk, N., & Olsen, C. E. (2012). Glucosinolate structures in evolution. Phytochemistry, 77, 16–45. doi:10.1016/j.phytochem.2012.02.005

Beilstein, M. A., Al-Shehbaz, I. A., Mathews, S., & Kellogg, E. A. (2008). Brassicaceae phylogeny inferred from phytochrome A and ndhF sequence data: tribes and trichomes revisited. American Journal of Botany, 95(10), 1307–1327. doi:10.3732/ajb.0800065

Benson, J., Pasquale, A., Van Driesche, R., & Elkinton, J. (2003). Assessment of risk posed by introduced braconid wasps to *Pieris virginiensis*, a native woodland butterfly in New England. Biological Control, 26(1), 83–93. doi:10.1016/S1049-9644(02)00119-6

Berenbaum, M. R., Favret, C., & Schuler, M. A. (1996). On defining “key innovations” in an adaptive radiation: Cytochrome P450s and Papilionidae. The American Naturalist, 148, 139–155. doi:10.1086/285907

Bolger, A. M., Lohse, M., & Usadel, B. (2014). Trimmomatic: A flexible trimmer for Illumina sequence data. Bioinformatics, 30(15), 2114–2120. doi:10.1093/bioinformatics/btu170

Bolter, C., & Jongsma, M. A. (1997). The adaptation of insects to plant protease inhibitors. Journal of Insect Physiology, 43(10), 885–895.

Bond, J. E., & Opell, B. D. (1988). Testing adaptive radiation and key innovation hypotheses in spiders. Evolution, 52(2), 403–414.

Broadway, R. M. (1989). Characterization and ecological implications of midgut proteolytic activity in Larval Pieris rapae and Trichoplusia ni. Journal of Chemical Ecology, 15(7), 2102–2113.

Broadway, R. M. (1996). Dietary proteinase inhibitors alter complement of midgut proteases. Archives of Insect Biochemistry and Physiology, 32(1), 39–53.

Camacho, C., Coulouris, G., Avagyan, V., Ma, N., Papadopoulos, J., Bealer, K., & Madden, T. L. (2009). BLAST+: Architecture and applications. BMC Bioinformatics, 10, 1–9. doi:10.1186/1471-2105-10-421

Capella-Gutiérrez, S., Silla-Martínez, J. M., & Gabaldón, T. (2009). trimAl: a tool for automated alignment trimming in large-scale phylogenetic analyses. Bioinformatics (Oxford, England), 25(15), 1972–1973. doi:10.1093/bioinformatics/btp348

Cock, P. J. A., Chilton, J. M., Grüning, B., Johnson, J. E., & Soranzo, N. (2015). NCBI BLAST+ integrated into Galaxy. GigaScience, 4(39). doi:10.1186/s13742-015-0080-7

Couvreur, T. L. P., Franzke, A., Al-Shehbaz, I. A., Bakker, F. T., Koch, M. A., & Mummenhoff, K. (2010). Molecular phylogenetics, temporal diversification, and principles of evolution in the mustard family (Brassicaceae). Molecular Biology and Evolution, 27(1), 55–71. doi:10.1093/molbev/msp202

Edger, P. P., Heidel-Fischer, H. M., Bekaert, M., Rota, J., Glöckner, G., Platts, A. E., … Wheat, C. W. (2015). The butterfly plant arms-race escalated by gene and genome duplications. Proceedings of the National Academy of Sciences of the United States of America, 112, 8362–8366. doi:10.1073/pnas.1503926112

Fahey, J. W., Zalcmann, A. T., & Talalay, P. (2001). The chemical diversity and distribution of glucosinolates and isothiocyanates among plants. Phytochemistry, 56(1), 5–51.

Fischer, H. M., Wheat, C. W., Heckel, D. G., & Vogel, H. (2008). Evolutionary origins of a novel host plant detoxification gene in butterflies. Molecular Biology and Evolution, 25(5), 809–820. doi:10.1093/molbev/msn014

Franzke, A., Lysak, M. A., Al-Shehbaz, I. A., Koch, M. A., & Mummenhoff, K. (2011). Cabbage family affairs: the evolutionary history of Brassicaceae. Trends in Plant Science, 16(2), 108–116. doi:10.1016/j.tplants.2010.11.005

Götz, S., García-Gómez, J. M., Terol, J., Williams, T. D., Nagaraj, S. H., Nueda, M. J., … Conesa, A. (2008). High-throughput functional annotation and data mining with the Blast2GO suite. Nucleic Acids Research, 36(10), 3420–3435. doi:10.1093/nar/gkn176

Grabherr, M. G., Haas, B. J., Yassour, M., Levin, J. Z., Thompson, D. A., Amit, I., … Regev, A. (2011). Full-length transcriptome assembly from RNA-Seq data without a reference genome. Nature Biotechnology, 29(7), 644–652. doi:10.1038/nbt.1883

Harvey, J. A., Poelman, E. H., & Gols, R. (2010). Development and host utilization in Hyposoter ebeninus (Hymenoptera: Ichneumonidae), a solitary endoparasitoid of *Pieris rapae* and *P. brassicae* caterpillars (Lepidoptera: Pieridae). Biological Control, 53(3), 312–318. doi:10.1016/j.biocontrol.2010.02.004

Heidel-Fischer, H. M., Vogel, H., Heckel, D. G., & Wheat, C. W. (2010). Microevolutionary dynamics of a macroevolutionary key innovation in a Lepidopteran herbivore. BMC Evolutionary Biology, 10, 60. doi:10.1186/1471-2148-10-60

Hoang, D. T., Chernomor, O., Von Haeseler, A., Minh, B. Q., & Vinh, L. S. (2018). UFBoot2: Improving the ultrafast bootstrap approximation. Molecular Biology and Evolution, 35(2), 518–522. doi:10.1093/molbev/msx281

Hunter, J. P. (1998). Key innovation and the ecology of macroevolution. Trends in Ecology & Evolution, 13(1), 31–36.

Janz, N. (2011). Ehrlich and Raven revisited: mechanisms underlying codiversification of plants and enemies. Annual Review of Ecology, Evolution, and Systematics, 42(1), 71–89. doi:10.1146/annurev-ecolsys-102710-145024

Kitahara, H. (2016). Oviposition plants and seasonal migratory movements of sympatric *Pieris melete* and *P. napi japonica* (Lepidoptera, Pieridae). Lepidoptera Science, 67(1), 32–40.

Kosakovsky Pond, S. L., Frost, S. D. W., & Muse, S. V. (2005). HyPhy: Hypothesis testing using phylogenies. Bioinformatics, 21(5), 676–679. doi:10.1093/bioinformatics/bti079

Loytynoja, A., & Goldman, N. (2005). An algorithm for progressive multiple alignment of sequences with insertions. Proceedings of the National Academy of Sciences, 102(30), 10557–10562. doi:10.1073/pnas.0409137102

Nei, M., & Gojoborit, T. (1986). Simple methods for estimating the numbers of synonymous and nonsynonymous nucleotide substitutions. Molecular Biology and Evolution, 3(5), 418–426. doi:10.1093/oxfordjournals.molbev.a040410

Nguyen, L. T., Schmidt, H. A., Von Haeseler, A., & Minh, B. Q. (2015). IQ-TREE: A fast and effective stochastic algorithm for estimating maximum-likelihood phylogenies. Molecular Biology and Evolution, 32(1), 268–274. doi:10.1093/molbev/msu300

Ohsaki, N., & Sato, Y. (1994). Food plant choice of *Pieris* butterflies as a trade-off between parasitoid avoidance and quality of plants. Ecology, 75(1), 59–68. doi:10.2307/1939382

Olsen, C. E., Huang, X. C., Hansen, C. I. C., Cipollini, D., Ørgaard, M., Matthes, A., … Agerbirk, N. (2016). Glucosinolate diversity within a phylogenetic framework of the tribe Cardamineae (Brassicaceae) unraveled with HPLC-MS/MS and NMR-based analytical distinction of 70 desulfoglucosinolates. Phytochemistry, 132, 33–56. doi:10.1016/j.phytochem.2016.09.013

Randall, T. A., Perera, L., London, R. E., & Mueller, G. A. (2013). Genomic, RNAseq, and molecular modeling evidence suggests that the major allergen domain in insects evolved from a homodimeric origin. Genome Biology and Evolution, 5(12), 2344–2358. doi:10.1093/gbe/evt182

RStudioTeam. (2016). RStudio: Integrated Development for R. Retrieved from http://www.rstudio.com

Schweizer, F., Heidel-Fischer, H., Vogel, H., & Reymond, P. (2017). *Arabidopsis* glucosinolates trigger a contrasting transcriptomic response in a generalist and a specialist herbivore. Insect Biochemistry and Molecular Biology, 85, 21–31. doi:10.1016/j.ibmb.2017.04.004

Simon, J.-C., d’Alencon, E., Guy, E., Jacquin-Joly, E., Jaquiery, J., Nouhaud, P., … Streiff, R. (2015). Genomics of adaptation to host-plants in herbivorous insects. Briefings in Functional Genomics, 14(6), 413–423. doi:10.1093/bfgp/elv015

Smith, M. D., Wertheim, J. O., Weaver, S., Murrell, B., Scheffler, K., & Kosakovsky Pond, S. L. (2015). Less is more: An adaptive branch-site random effects model for efficient detection of episodic diversifying selection. Molecular Biology and Evolution, 32(5), 1342–1353. doi:10.1093/molbev/msv022

Stamatakis, A. (2014). RAxML version 8: A tool for phylogenetic analysis and post-analysis of large phylogenies. Bioinformatics, 30(9), 1312–1313. doi:10.1093/bioinformatics/btu033

Tibshirani, R., Walther, G., & Hastie, T. (2001). Estimating the number of clusters in a data set via the gap statistic. Journal of the Royal Statistical Society. Series B: Statistical Methodology, 63(2), 411–423. doi:10.1016/j.scico.2012.08.004

Ueno, M. (1997). A note on the large white, *Pieris brassicae*.(I). Yadoriga, 169, 25–41.

Weaver, S., Shank, S. D., Spielman, S. J., Li, M., Muse, S. V, & Kosakovsky Pond, S. L. (2018). Datamonkey 2.0: A modern web application for characterizing selective and other evolutionary processes. Molecular Biology and Evolution, 35(3), 773–777. doi:10.1093/molbev/msx335

Wheat, C. W., Vogel, H., Wittstock, U., Braby, M. F., Underwood, D., & Mitchell-Olds, T. (2007). The genetic basis of a plant-insect coevolutionary key innovation. Proceedings of the National Academy of Sciences of the United States of America, 104(51), 20427–20431. doi:10.1073/pnas.0706229104

Wittstock, U., Agerbirk, N., Stauber, E. J., Olsen, C. E., Hippler, M., Mitchell-Olds, T., … Vogel, H. (2004). Successful herbivore attack due to metabolic diversion of a plant chemical defense. Proceedings of the National Academy of Sciences of the United States of America, 101(14), 4859–4864. doi:10.1073/pnas.0308007101

Yang, Z. (2007). PAML 4: Phylogenetic analysis by maximum likelihood. Molecular Biology and Evolution, 24(8), 1586–1591. doi:10.1093/molbev/msm088

