## Supplementary information for "Molecular signatures of selection associated with host-plant differences in *Pieris* butterflies"

**Table S1**

**25 Brassicaceae plants used for the feeding assay**

| Plant species | Seed collection site |
| --- | --- |
| *Arabidopsis kamchatica* | Nagano, Japan (36° 31' N, 138° 20' E) |
| *Arabidopsis thaliana* | Hokkaido, Japan (42° 50' N, 141° 41' E) |
| *Arabis hirsuta* | Nagano, Japan (36° 31' N, 138° 20' E) |
| *Aubrieta deltoidea* | - |
| *Aurinia saxatilis* | - |
| *Barbarea orthoceras* | Toyama, Japan (36° 34' N, 137° 26' E) |
| *Berteroa incana* | Burlington, Canada (43° 30' N, 79° 79' W) |
| *Brassica napus* | Chiba, Japan (35° 30' N, 140° 50' E) |
| *Brassica tournefortii* | Chiba, Japan (35° 54' N, 139° 56' E) |
| *Capsella bursa-pastoris* | Chiba, Japan (35° 30' N, 140° 50' E) |
| *Cardamine hirsuta* | Chiba, Japan (35° 30' N, 140° 50' E) |
| *Cardamine occulta* | Chiba, Japan (35° 30' N, 140° 50' E) |
| *Descurainia sophia* | Inner Mongolia, China (43° 37' N, 116° 42' E) |
| *Draba nemorosa* | Nagano, Japan (36° 20' N, 137° 50' E) |
| *Eruca sativa* | - |
| *Eruca sylvatica* | - |
| *Erysimum cheiranthoides* | Hokkaido, Japan (42° 50' N, 141° 41' E) |
| *Lepidium sativum* | - |
| *Matthiola longipetala* | - |
| *Nasturtium officinale* | Chiba, Japan (35° 30' N, 140° 50' E) |
| *Raphanus sativus var. raphanistroides* | Chiba, Japan (35° 07' N, 140° 11' E) |
| *Rorippa indica* | Chiba, Japan (35° 30' N, 140° 50' E) |
| *Sisymbrium orientale* | Chiba, Japan (35° 54' N, 139° 56' E) |
| *Thlaspi arvense* | Hokkaido, Japan (42° 56' N, 142° 01' E) |
| *Turritis glabra* | Nagano, Japan (36° 31' N, 138° 20' E) |
